# Algorithms underlying flexible phototaxis in larval zebrafish

**DOI:** 10.1101/2020.07.18.210260

**Authors:** Alex B. Chen, Diptodip Deb, Armin Bahl, Florian Engert

## Abstract

To thrive, organisms must maintain physiological and environmental variables in optimal ranges. However, in a dynamic world, the optimal range of a variable might fluctuate depending on the organism’s state or environmental conditions. Given these fluctuations, how do biological control systems maintain optimal control of physiological and environmental variables? We explored this question by studying the phototactic behavior of larval zebrafish. We demonstrate, with behavioral experiments and computational modeling, that larval zebrafish use phototaxis to maintain environmental luminance at a set point that depends on luminance history. We further show that fish compute this set point using information from both eyes, and that the set point fluctuates on a timescale of seconds when environmental luminance changes. These results expand on previous studies, where phototaxis was found to be primarily positive, and suggest that larval zebrafish, rather than consistently turning towards the brighter areas, exert homeostatic control over the luminance of their surroundings. Furthermore, we show that fluctuations in the surrounding luminance feed back on the system to drive allostatic changes to the luminance set point. Our work has uncovered a novel principle underlying phototaxis in larval zebrafish and characterized a behavioral algorithm by which larval zebrafish exert control over a sensory variable.

## INTRODUCTION

All living organisms exert control over a variety of physiological variables. For example, animals employ sophisticated control systems to keep body temperature, body weight, blood osmolarity, as well as many other parameters, within narrow ranges critical for bodily function [1]. Many of these homeostatic processes involve comparing moment-to-moment values of the controlled physiological variables to ‘set points’ that the control system seeks to maintain. When a variable deviates from its set point, the control system acts, often via negative feedback, to restore the variable to its set value.

Conceptualizations of homeostatic control often treat set points as fixed in value, but changing environmental or internal conditions could render an existing, fixed, set point maladaptive [2]. When this occurs, a robust control system ought to flexibly adjust its set point to a range adaptive to the new conditions. This process has been termed *allostasis* [3,4], and allostatic shifts in set points occur everywhere across the animal kingdom. For example, many endothermic animals exhibit an elevated body temperature set point, fever, in response to infection [5,6], whereas animals that hibernate through the winter reduce their body temperature set point during hibernation, but increase their caloric set point before hibernation sets in [7,8]. Finally, homeostatic and allostatic control can involve behavioral, in addition to physiological, changes. Ectothermic animals that regulate body temperature by seeking out warmer or cooler regions of the environment also exhibit behavioral fever [9]. Despite the ubiquity of allostasis in physiology, it is still poorly understood how physiological control systems adjust their set points in response to changing internal and external conditions, and how allostasis interacts with homeostatic control.

In this study, we establish luminance-based navigation in larval zebrafish as a model for investigating behavioral allostatic control. Previous work on luminance-based navigation in larval zebrafish has focused on their tendency to orient and swim towards brighter regions of luminance gradients; this behavior is termed *positive phototaxis* [10–15]. However, in naturalistic environments, luminance varies widely, both throughout the day and as fish move into and out of shade. Therefore, a strategy of purely positive phototaxis might not be adaptive to larval zebrafish, and it is likely too simplistic a view of this complex behavior. Indeed, evidence of flexibility in the phototactic behavior of larval zebrafish has been documented. One study demonstrated that larval zebrafish avoid light sources that are too bright and that this avoidance depends on the luminance to which fish are preadapted [16]. Another study revealed that larval zebrafish exhibit negative phototaxis in gradients of near-infrared light [17]. These findings suggest that the phototactic behavior of larval zebrafish is flexible and can be modulated by environmental luminance and its history.

We sought to characterize the behavioral algorithms that underlie the flexibility of phototaxis in larval zebrafish. Towards that end, we delivered closed-loop luminance gradient stimuli to freely swimming larval zebrafish that were preadapted to different luminance histories, and we formulated simple behavioral algorithms that could explain the resultant behavior. We found that larval zebrafish perform positive and negative phototaxis to orient towards a set point luminance, the value of which depends on the luminance history of their surroundings. Furthermore, fish compute the set point luminance using visual information from both eyes. These findings uncovered previously unappreciated principles underlying phototaxis in larval zebrafish, namely that the larval zebrafish employs phototaxis to maintain its experienced luminance at a fixed value and that its luminance preference fluctuates in response to changing environmental luminance conditions. We believe that this behavioral control of experienced luminance can serve as a model for investigating neural implementations of homeostatic and allostatic control.

## RESULTS

### Larval zebrafish orient towards a set point during luminance-based navigation

To deliver controlled luminance stimuli to freely swimming larval zebrafish, we employed a closed-loop video projection system (Figure 1A) used in previous work [18] (STAR Methods). Here, a high-speed camera recorded a video of a fish swimming in a shallow, circular dish, a computer-vision program then calculated its position and orientation in real time, and a projector used this information to deliver visual stimuli fixed to the fish’s reference frame. As a result, the visual stimuli, and in particular a specific luminance, can be kept constant on the fish’s eyes, even if the animal moves continuously through the arena (see Methods).

**Figure 1.**
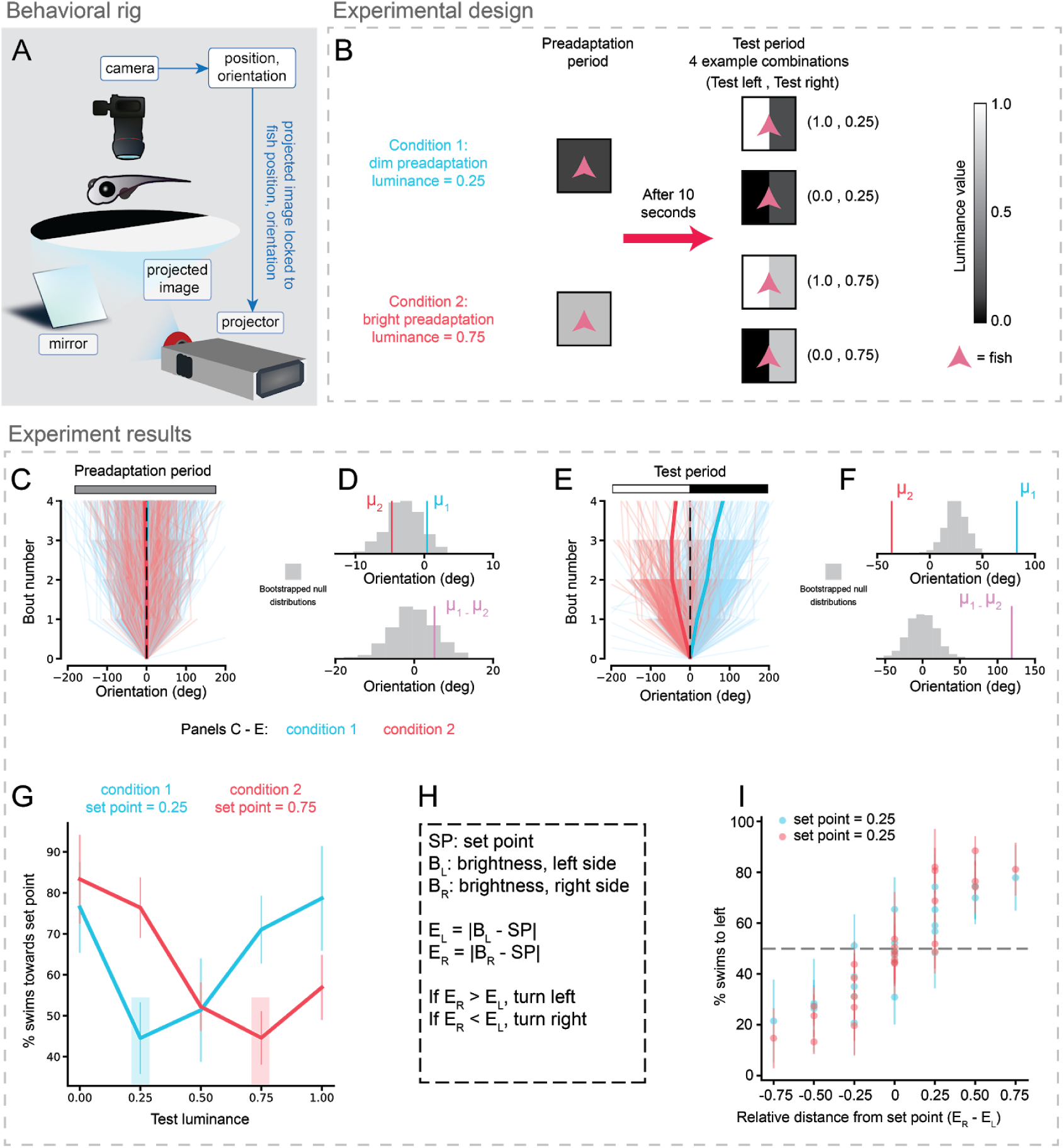
Larval zebrafish orient towards a set point during luminance-based navigation. **A**. Schematic of experimental setup. Larval zebrafish swim freely while visual stimuli are presented locked to fish reference frames (see STAR Methods). **B**. Experimental design. Each trial consisted of two periods. During the first period (‘preadaptation’) fish were held in a dim (L = 0.25, n = 11 fish) or bright (L = 0.75, n = 8 fish) environment for at least 10 seconds. Immediately following preadaptation, fish were subjected to a test period for at least 3 seconds. During the test period, the left and right sides of the environment relative to the fish were held fixed at brightness values (B_L_, B_R_), where B_L_ and B_R_ were selected pseudo-randomly from the set {0, 0.25, 0.50, 0.75, 1.00}. **C**. Change in orientation of fish over the last 3 s of preadaptation. Color bar reflects equal luminance on both sides of the fish. Each trace shows the change in orientation in a single trial for a single animal. Orientation values are presented relative to the animal’s orientation at 3 s before test period onset. Blue traces: condition 1 (11 fish, 2200 trials). Red traces: condition 2 (8 fish, 720 trials). Thicker traces show the mean of each condition (partially overlapping with dotted line). **D**. Statistical comparison between preadaptation periods in conditions 1 and 2. Top: gray histogram shows bootstrapped distribution of trials shuffled randomly and split with 2200 trials in one group and 720 in another to preserve group size (1000 bootstrapped means). Red and blue ticks show observed means for conditions 1 and 2 respectively (μ1 = 0.40 deg, μ2 = -4.72 deg); they fall within the 90% CI of shuffled mean, [-7.31 deg, 1.26 deg]. Bottom histogram is the bootstrapped null distribution of the difference in means between the two conditions (1000 bootstrapped differences). Trials were shuffled and sorted as described above. Purple tick shows the observed difference in means (μ1 - μ2 = 5.12 deg), which falls within the 80% CI of the shuffled difference, [-7.25 deg, 6.89 deg]. **E**. Same as C for test period. Negative orientation angles were defined to be in the bright direction and positive orientation angles were defined to be in dim direction, as shown in the color bar. Note clear separation between the two conditions. **F**. Same analyses as D for test period. Observed means and difference between means fall completely outside bootstrapped distributions for shuffled data (1000 bootstrapped values for each distribution), indicating difference significant to p < 0.001. **G**. Analysis of the test period for trials in which one side of the fish was set point luminance. The luminance of the other side was considered the ‘test luminance’. Shaded region denotes set point luminance. Bias towards the set point side depended significantly on the test luminance (ANOVA, df = 4, p < 0.05 for both conditions). Bias towards set point luminance was higher when the test luminance deviated from set point than when test luminance was equal to set point luminance (condition 1: test luminance 0.25 vs. test luminance 1.0, t-test p < 0.01; condition 2: test luminance 0.75 vs. test luminance 0.0, t-test p < 0.01). Error bars denote standard deviation across fish. **H**. Explanation of abbreviations and calculations used to generate Figure 1I. **I**. Percent of leftward swims plotted as a function of E_R_ - E_L_. Higher E_R_ - E_L_ values drive higher leftward swim bias (t-test on slope of linear fit, p < 0.001).

Larval zebrafish were preadapted to either a bright (L = 0.75, see Methods), or a dim (L = 0.25) luminance level for 10 seconds (Figure 1B, preadaptation period). Following this preadaptation period, the fish experienced a split luminance environment, a test period, in which one visual hemifield was bright (L = 1) and the other visual hemifield was dim (L = 0) (Figure 1B). As shown in previous work [19–24], during the preadaptation period, when fish experienced uniform luminance, we observed no significant bias in turn direction (Figure 1C-D).

However, a comparison of the swimming statistics during the preadaptation period (Figure 1C-D) and the test period (Figure 1E-F) revealed that fish do not simply perform positive phototaxis, as has been suggested by previous studies [10,11,13,14], but that they consistently turn towards the side closest in luminance to the preadaptation period. This indicates that larval zebrafish prefer a luminance similar to the preadaptation luminance, suggesting that this value serves as a set point for subsequent luminance-driven navigation. Thus, fish preadapted to a bright environment exhibited a turning bias towards the bright visual hemifield during the test period, while fish preadapted to a dim environment exhibited a turning bias towards the dim visual hemifield during the test period (Figure 1E-F). The turning bias was highest immediately after switching from the preadaptation period to the test period, and gradually declined throughout the test period (Supplemental Figure 1C-D). To limit the effect of this adaptation on our analyses, we considered turning behavior only within the first three seconds of the test period for the analyses presented in Figure 1.

If the preadaptation luminance is truly the luminance set point during the test period, then fish should prefer the preadaptation luminance to any other luminance. To investigate this, we held one visual hemifield constant at the preadaptation luminance during the test period and changed the luminance of the other hemifield over a large range of luminances (L = 0, 0.25, 0.5, 0.75, and 1). To normalize differences in the number of swim bouts across different trials, we calculated the percent of leftward swims (defined to be negative angles) and rightward swims (positive angles) for each trial. This measure corresponded well with the accumulated angle measure used in Figure 1C-F (Supplemental Figure 1E). We next tested a large set of combinations of pre-adaptation and test values and found that, for all of these, the preadaptation luminance appears to act as a set point for luminance-based navigation during the test period (Figure 1G).

How might we formally characterize this relationship between preadaptation luminance and turning behavior? To that end, we hypothesized a simple behavioral algorithm that the fish might use: in the test period, the fish compares the brightness values experienced by the left eye (**B**_**L**_) and the right eye (**B**_**R**_) to the set point (**SP**) to generate two error signals (**E**_**L**_ **=** |**B**_**L**_ **-SP**|, **E**_**R**_ **=** |**B**_**R**_ **-SP**|). It then biases turning towards the direction of the smaller error. To determine how well our model describes light-seeking behavior, we analyzed turning data across all preadaptation and test period luminance conditions (2 preadaptation luminances x 5 left visual hemifield test luminances x 5 right visual hemifield test luminances = 50 total conditions, Supplemental Figure 1F-G). For each condition, we calculated **E**_**R**_ **-E**_**L**_ and plotted the percent of swims to the left as a function of **E**_**R**_ **-E**_**L**_ (Figure 1H-I). Consistent with our model’s predictions, when **E**_**R**_ was larger than **E**_**L**_ (**E**_**R**_ **-E**_**L**_ **> 0**), fish swam leftward, and when **E**_**R**_ was smaller than **E**_**L**_ (**E**_**R**_ **-E**_**L**_ **< 0**), fish swam rightward.

These results indicate that the larval zebrafish prefers the environmental luminance to which it is adapted. When it encounters environments of variable luminance, the fish moves to minimize deviations from the adapted luminance. We note that this behavior appears similar to the behavioral defense of a homeostatic set point, as seen in thermoregulation in larval zebrafish [25].

### The luminance set point is computed using luminance information from both eyes

While our hypothesized behavioral algorithm describes the relationship between preadaptation luminance and split-luminance turning bias well, it assumes a common set point to which luminance values from both the left and right eyes are compared. An alternative to this unitary, binocular set point is two separate set points – one for the left eye and another for the right eye. Determining whether two monocular set points exist (Figure 2A), or whether only one binocular set point exists (Figure 2B), would inform hypotheses about where this set point is implemented in the brain. If two monocular set points exist, they might be implemented in earlier regions of the visual processing stream, such as the retina, before information from the two eyes have converged. On the other hand, if only one binocular set point exists, it must be implemented in a brain region that integrates information from both eyes.

**Figure 2.**
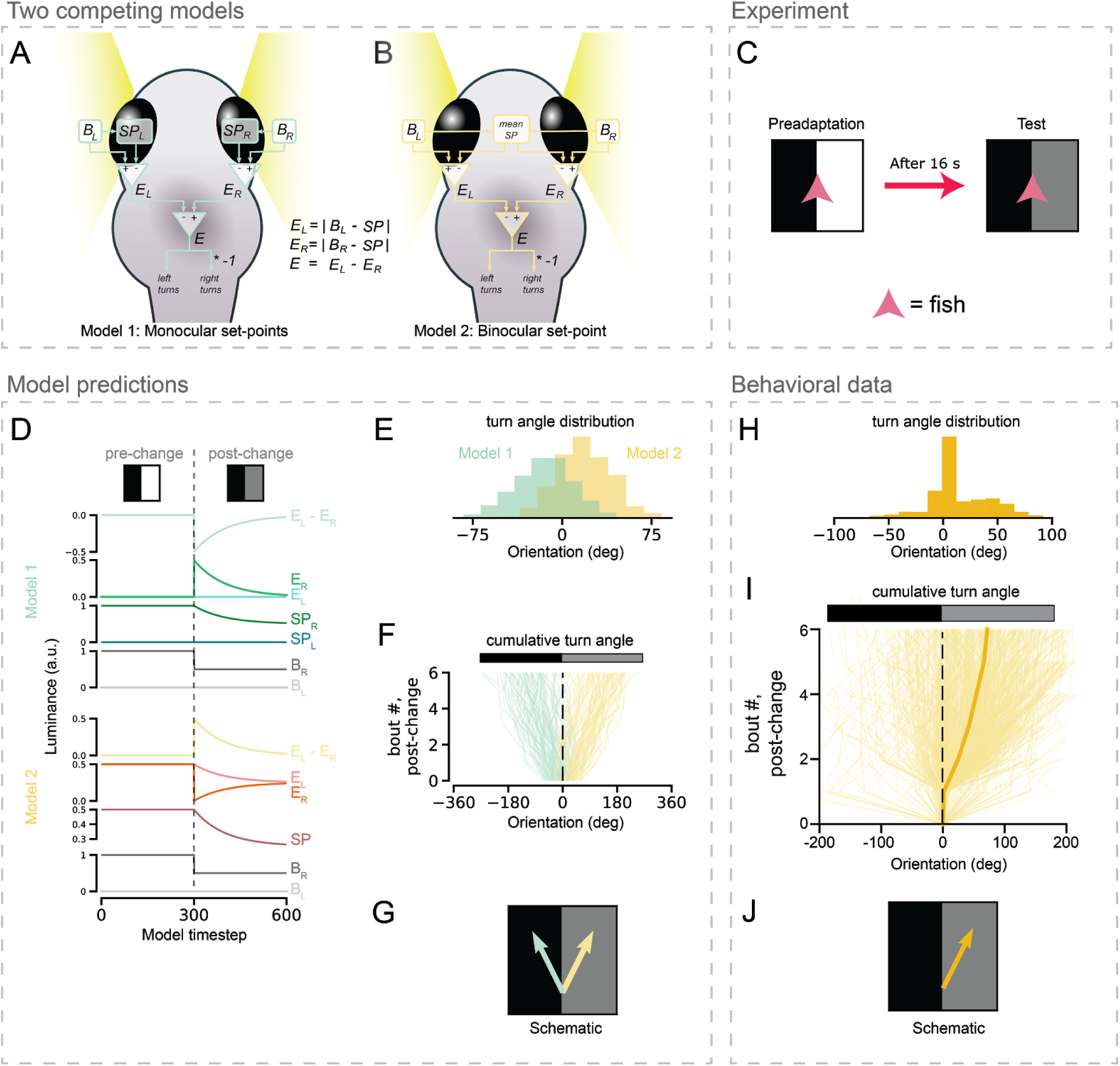
The luminance set point is computed using luminance information from both eyes. **A** and **B**. Schematics of two competing hypotheses for the behavioral algorithm that larval zebrafish use for set point seeking. **A**. Each eye has its own luminance set point (SP_L_ and SP_R_). B_L_: brightness experienced by left eye. B_R_: brightness experienced by right eye. **B**. Unitary set point (SP) that approaches the mean of B_R_ and B_L_. Other computations are identical to A except B_R_ and B_L_ are compared to SP instead of SP_L_ and SP_R_. See STAR Methods for implementation of models. **C**. Experimental setup. During preadaptation, one side of the fish was bright and the other side was dark. During the test period, the luminance of the bright side was decreased to a final value between two preadaptation values. **D**. Simulated evolution of parameters for Models 1 and 2 for experiment described in C. Note the opposite signs of E_L_ - E_R_ predicted by the two models during the test period. **E**. Simulated turn angle distribution of the two models. Means are significantly different (exact test, p < 0.05). **F**. Cumulative turn angle over the first 6 bouts predicted by two models. Color bar shows luminance; positive angles are defined to be towards higher luminance. **G**. Cartoon schematic showing that the two models predict opposite behavioral outcomes. **H**. Turn angle distribution and **I**. Cumulative turn angle observed in real fish. Mean turn angle and cumulative turn angle significantly greater than 0 (exact tests, p < 0.01), indicating consistency with model 2, schematized in **J**.

The experimental paradigm described in Figure 1B cannot differentiate between the two competing hypotheses schematized in Figure 2A-B because during the preadaptation period, the two eyes experience the same luminance conditions, so we cannot tell whether preadaptation generates a unitary, binocular set point, or two separate monocular set points with the same luminance value. Therefore, we performed experiments in which we preadapted the fish to a split-luminance environment, in which one visual hemifield was relatively bright (L = 1) and the other visual hemifield was relatively dim (L = 0) for 16 seconds (Figure 2C). Following this preadaptation period, we changed the luminance of either the bright (Figure 2C) or dim (Supplemental Figure 2E) visual hemifield to an intermediate value (L = 0.5), while keeping the other one constant.

To show the differences in behavior predicted by these two competing models, we implemented both models computationally and simulated behavioral outcomes of the experiment presented in Figure 2C (see STAR Methods for details of model implementation). If each eye possesses its own luminance set point, then changing the luminance experienced by one eye would generate an error signal (Figure 2D) that drives a turning bias away from that direction (Figure 2E-F, green). On the other hand, if luminance-based navigation is driven by a single set point, i.e. the mean luminance value of the two eyes (L = 0.5), then the fish should turn away from the eye receiving constant illumination (Figure 2E-F, yellow), since there the error signal remains high (Figure 2D, E_L_). In the other eye the error drops from a high value to zero (Figure 2D, E_R_).

We next tested these predictions from our model in behavioral experiments (Fig 2G-I). We observed that when the bright side of the split-luminance environment was dimmed to an intermediate value, the fish exhibited a turning bias towards the changed side and away from the constant side (Figure 2H, Supplemental Figure 2 C-D). This turning bias does not exist during the preadaptation period (Supplemental Figure 2 A-B). Finally, we also observed a turning bias, relative to preadaptation, when the dim preadaptation side was brightened (Supplemental Figure 2 E). While this effect was more subtle than when the bright preadaptation side was dimmed (Supplemental Figure 2 F-G), it was still statistically significant. A potential reason for the weaker effect upon brightening of the dim side might be that the measured luminance values are not translated linearly by the fish’s nervous system.

Nonetheless, these data are consistent with the existence of a unitary set point generated by averaging luminance values across both eyes.

### The luminance set point depends on environmental luminance history

The luminance of an animal’s surroundings changes over short timescales, as the animal moves into and out of shade, and over long timescales, with the rising and setting of the sun. If the luminance that a larval zebrafish seeks depends on the luminance to which the fish has been preadapted, then this set point luminance should change over equivalent time scales. This could explain the gradual decline in turning bias that we observe (Supplemental Figure 1C-D) in the seconds after the fish’s environment switches in luminance.

To more directly test our hypothesis that changing environmental luminance would alter the luminance set point of larval zebrafish, we performed experiments outlined in Figure 3A. First, we subjected fish to an initial preadaptation period, termed preadaptation 1 in either bright (L = 0.75) or dim (L = 0.25), uniform luminance environments (Figure 3A). Following preadaptation 1, we either transitioned the fish directly to a split-luminance test environment to assay turning bias (Figure 3A, test), or we instead transitioned the fish to a second preadaptation period (termed preadaptation 2) of variable length (Figure 3A). If preadaptation 1 was dim, then preadaptation 2 was bright (Figure 3A, condition 1), and vice versa (Figure 3A, condition 2).

**Figure 3.**
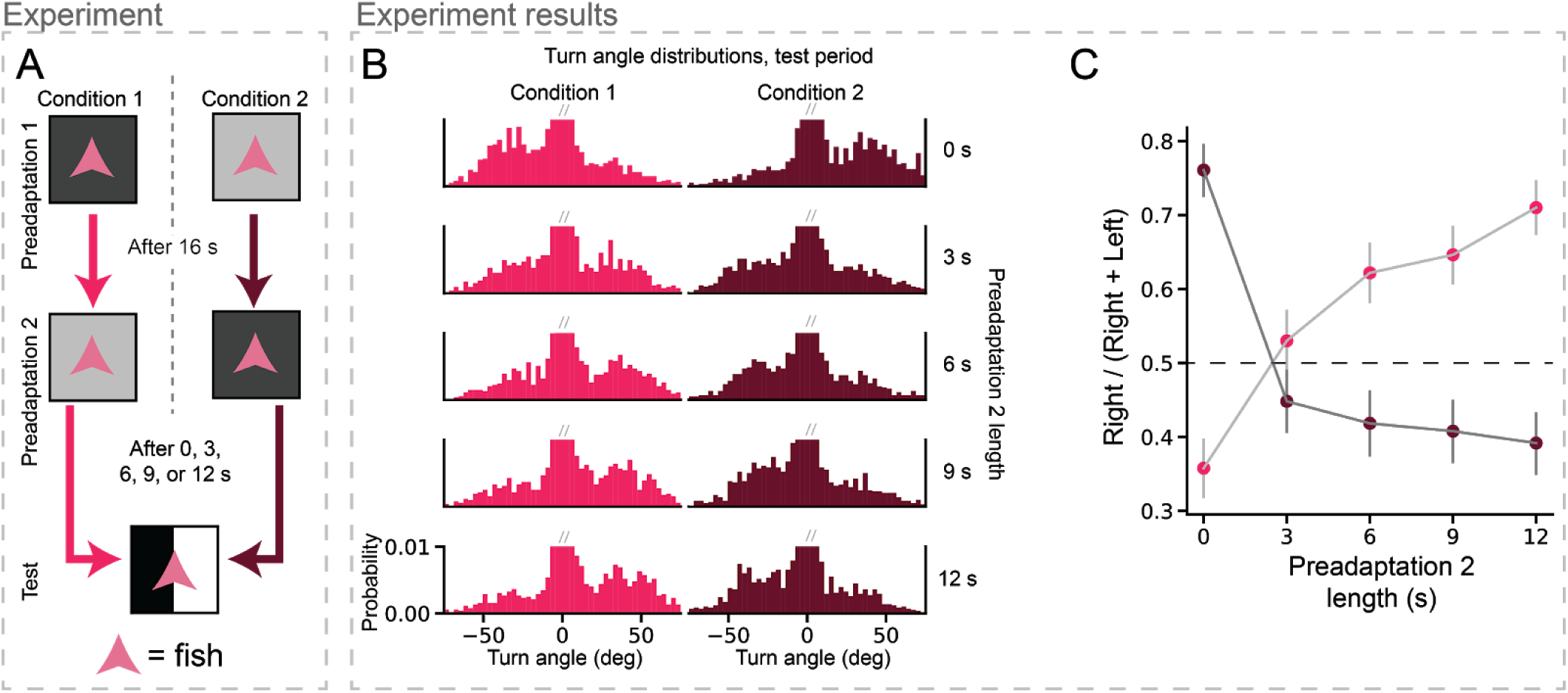
The luminance set point depends on environmental luminance history. **A**. Schematic of experiment to probe temporal evolution of set point. Fish are subjected to a first preadaptation period in either dim (L = 0.25, condition 1) or bright (L = 0.75, condition 2) luminance. Fish then experience a second preadaptation period of varying length (condition 1: luminance increases to 0.75, condition 2: luminance decreases to 0.25). Finally, fish experience a split luminance test period (L = 0 on left, 1 on right). **B**. Turn angle probability distributions for both conditions and different preadaptation 2 lengths. Low-angle swim bins are truncated (gray slashes) to allow for comparison of changes to larger-angle turn distributions. N = 14 fish, **C**. Number of right turns as a fraction of total turns. Angle threshold for turn classification was 15 degrees. Note opposite turning bias for 0 s preadaptation 2 and 12 s preadaptation 2 in both conditions (t-test, p < 0.001 for both conditions).

When fish transitioned from preadaptation 1 directly into the test environment (i.e. 0 s of preadaptation 2), fish exhibited a turning bias towards the preadaptation 1 luminance (Figure 3B-C, 0 s preadaptation 2), consistent with our findings presented in Figure 1. Strikingly, we observe an asymmetry in the maximal preferences driven by light versus dark preadaptation. Light preadaptation drove a stronger preference for light than dark preadaptation drove for darkness (Figure 3C). On the other hand, if we held the fish in preadaptation 2, the fish instead exhibited a turning bias towards the latter luminance (Figure 3B-C). This switch in turning bias due to the second preadaptation period varied with its duration; a longer preadaptation drove a larger change in luminance set point (Figure 3C). By contrast, when the second preadaptation was relatively short, around 3s in duration, fish exhibited no strong bias towards either luminance during the split luminance period. We conclude that environmental luminance fluctuations drive allostatic changes in the luminance set point over a timescale of seconds.

Taken with our observations that larval zebrafish seek a luminance set point, these data suggest that luminance-based navigation in larval zebrafish can be described by a ‘homeostatic-allostatic’ model. Over short timescales, the larval zebrafish exerts control over the luminance it experiences by using positive and negative phototaxis to orient towards a luminance set point. However, when the environmental luminance changes, the fish’s luminance set point is allostatically modulated to reflect the new mean environmental luminance. We speculate that this allostatic modulation of the fish’s set point might be coordinated with physiological changes in the fish’s visual system [26] that adapts it to the new environmental luminance.

### Luminance set point seeking is consistent with previously reported positive phototactic behavior

Why have there been many robust observations of positive phototaxis in larval zebrafish but little previous evidence of the luminance set point seeking that we report here? One possibility is that fish have a stronger maximal preference towards light than towards darkness as seen in Figure 3C. In addition, we argue that because these previous studies generally preadapted fish to an environment brighter than the test environment, the fish showed a preference for brighter regions during the test period. Consequently, fish would orient and swim towards brighter regions of the test environment and thus exhibit positive phototaxis. Indeed, when previous studies preadapted fish to environments darker than the test environment, fish exhibited much less positive [14], and sometimes even negative, phototaxis [16].

To demonstrate that an agent using our proposed homeostatic-allostatic algorithm for luminance-based navigation would perform positive phototaxis in similar experimental conditions used by previous studies, we modeled the behavior of virtual fish employing the algorithm schematized in Figure 2B. This algorithm is implemented by comparing the luminances of the left and right eyes to a common set point, and biasing swims towards the side closer to the set point. Furthermore, the set point evolves over time to approach the mean luminance of the two eyes (STAR Methods). Note that our goal was not to recapitulate our and previous data with quantitative precision but instead to show qualitatively that the same agent can perform set point seeking and positive phototaxis without fine-tuning of parameters.

Using the homeostatic-allostatic model, we first sought to replicate the history-dependent, set point seeking behavior reported in Figure 1C. We subjected model fish to the experimental conditions outlined in Figure 1B (Figure 4A). Consistent with our reported results from Figure 1, fish preadapted in the bright environment preferred the bright side of the test environment, while fish preadapted to the dim environment preferred the dim side (Figure 4B). These results were qualitatively similar to those reported in Figure 1B (Figure 4C).

**Figure 4.**
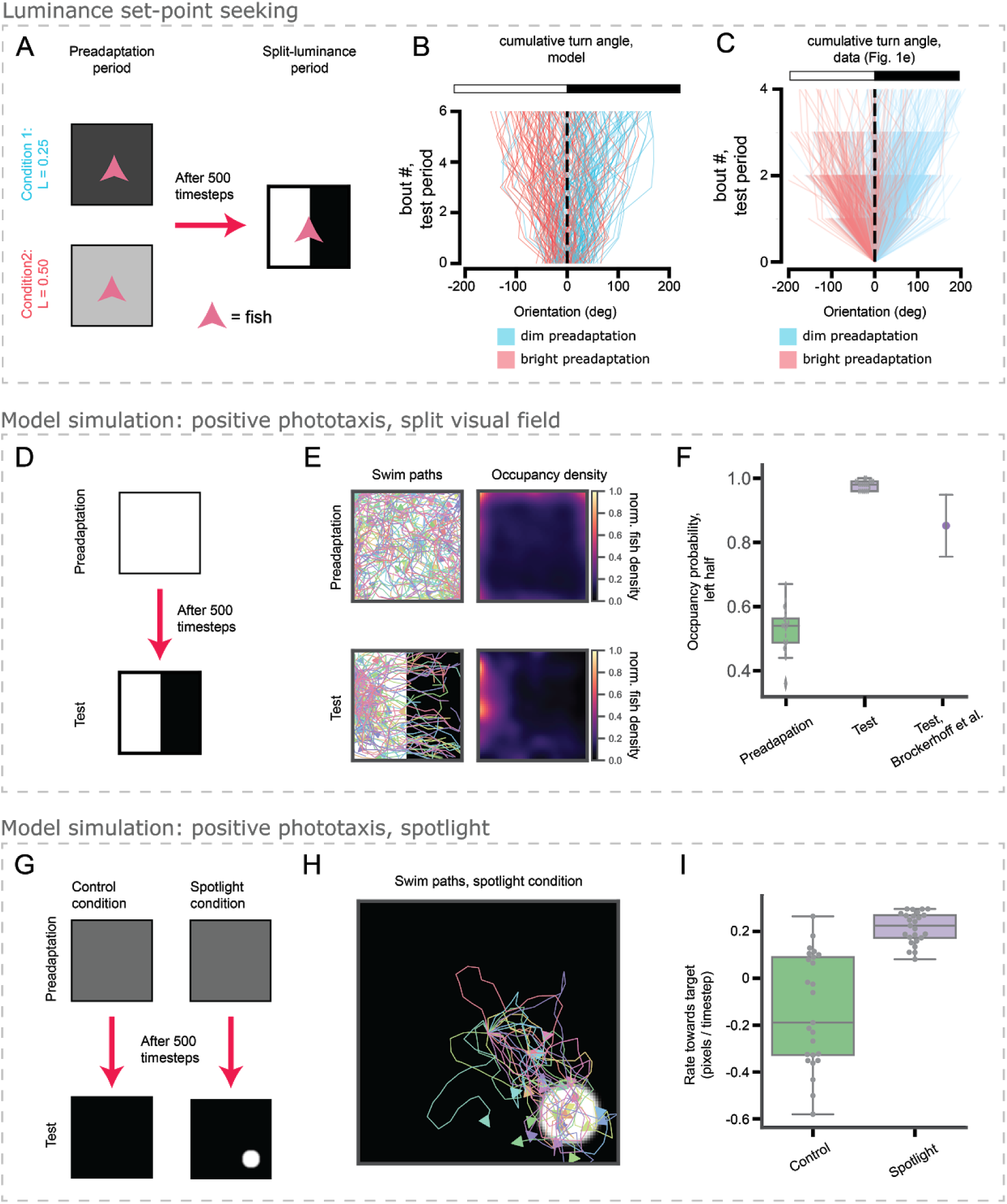
Luminance set point seeking is consistent with previously reported positive phototactic behavior. **A**. Schematic of experiment described in Figure 1B. **B**. Simulated fish results for experiment shown in A. Different conditions show opposite luminance preferences (exact test on difference between mean orientation after 6 bouts, p < 0.01). **C**. Replotted data from Figure 1C for comparison. **D-F**. Simulations of split-arena phototaxis experiment performed by [14]. **D**. Simulated fish swim freely in a brightly lit (L = 1) arena; after 500 simulation timesteps, the right half of the arena is darkened (L = 0). Note that the visual stimuli here are fixed in the lab reference frame, not the fish’s reference frame. **E**. Swim paths and occupancy densities of 100 simulated fish during preadaptation and test. Arrowheads denote final positions of the fish. **F**. Quantification of occupancy density of the left half of the arena during preadaptation and test periods for simulated fish and during the test period reported by [14]. Error bars denote standard deviation of mean occupancy. Preadaptation not significantly different from change (t-test, p > 0.25), Test significantly larger than chance (t-test, p < 0.001). **G**. Simulations of spotlight phototaxis experiment performed by Brockerhoff et al. [16]. In both conditions, fish are preadapted to a dim environment (L = 0.1). After 500 simulation timesteps, the environment either completely darkened (L = 0, control condition) or completely darkened (L = 0) save a spotlight (L = 1). The arena is 101 × 101 pixels, and the spotlight has a radius of 11 pixels. During preadaptation, fish positions were fixed to the middle of the arena. During the test period, fish were allowed to swim freely, and the visual scene was fixed to the lab reference frame, not the fish reference frame. **H**. Swim paths during the test period in the spotlight condition; fish swim towards the spotlight. **I**. Quantification showing that fish in the spotlight condition swim towards the spotlight more than fish in the control condition swim towards a fictive spotlight in the same location (t-test, p < 0.05, n = 25 fish in each condition).

Next, we sought to use the same model to replicate two different accounts of phototaxis in larval zebrafish (Figure 4D-I). In the first [14], model fish swam freely in a uniformly, brightly lit chamber for 500 model timesteps (Figure 4D, preadaptation); following this preadaptation period, we dimmed the right half of the chamber while keeping the left half bright (Figure 4D, test). Figure 4E exhibits the swim paths of 100 simulated fish, as well as their occupancy density in the chamber, during the preadaptation and test periods. The occupancy density in both conditions are quantified in Figure 4F and compared to the value reported by [14]. During the preadaptation periods, simulated fish showed no bias in occupancy between the left and right halves of the chamber (Figure 4F, preadaptation), a result expected due to symmetry. On the other hand, during the test period, fish exhibited a strong bias towards the brighter left side of the chamber (Figure 4F, test), as seen in [14].

We sought to replicate another observation of positive phototaxis made by Burgess et al. [16] using a smaller ‘spotlight’ of high luminance during the test period (Figure 4G). After preadapting simulated fish to an environment for 500 timesteps at a fixed location at the center of the chamber, fish experienced either a control or a spotlight environment. In the spotlight environment, we dimmed the chamber except for a small spotlight which was placed into the “target location” in the lower right. In the control environment, the entire chamber was dimmed. During this period, fish were free to swim around the chamber. We then let the simulation run for another 100 timesteps and measured the rate at which fish moved towards or away from the target location. In the control environment fish ended up, purely for geometrical reasons, further from the target (Figure 4I, control). However, in the spotlight environment, fish moved towards the target and thereby exhibited positive phototaxis (Figure 4I, spotlight).

We conclude from these data that our homeostatic-allostatic model of luminance-based navigation in larval zebrafish can generate both set point seeking behavior and previously observed positive phototaxis behavior in more realistic environments.

## DISCUSSION

Phototaxis occurs in many domains of life, from bacteria and unicellular eukaryotes [27,28] to plants [29,30] and animals [12,14,31–33]. While phototaxis benefits photosynthetic organisms by allowing them to find sources of energy [34], the purpose of phototaxis in heterotrophic animals that do not derive energy from light is not as obvious. Phototaxis likely serves a diverse set of functions across the animal kingdom; this diversity is reflected in the observed flexibility and context-dependence of phototactic behavior in animals like *Drosophila melanogaster*, honey bees, and sea anemones, [35–37]. Our work reveals that larval zebrafish may employ phototaxis in part to control the luminance levels of their surroundings. This principle unifies previous observations made about spatial phototaxis in larval zebrafish and helps to explain why environmental luminance affects phototactic strength and direction [14,16].

Notably, we observed that bright preadaptation could drive a larger turning bias than could dim preadaptation (Figure 3C). This asymmetry could explain why positive phototaxis is more often observed in larval zebrafish than negative phototaxis. It raises the possibility that larval zebrafish exhibit a residual positive phototactic behavior in addition to our observed luminance set point seeking. The mechanism underlying this asymmetry constitutes an interesting direction for future experiments and may yield insights on how luminance information is encoded by the larval zebrafish’s brain.

We speculate that control of environmental luminance benefits larval zebrafish because it enables them, over short timescales, to maintain luminance at a level to which their visual system is adapted. For example, if a larval zebrafish is dark-adapted, sudden brightening of its surroundings will initially overwhelm the dynamic range of its visual system and thereby degrade its contrast sensitivity. If the fish remains in the bright environment, light adaptation would eventually raise the dynamic range of visual processing to better suit its new environment. However, the physiological adaptation to changing luminance necessitates a shift of the animal’s luminance set point, or the animal would move towards the previous luminance to which it is no longer adapted. The short-term behavioral control of luminance we observe, and the longer-term shift in luminance preference, is similar to the allostatic modulation of a faster homeostatic control system. This motif, found in many homeostatic control systems, endows those systems with the ability to alter the value of their controlled variable to suit fluctuating environmental conditions.

Our characterization of the behavioral algorithm underlying luminance-based navigation in larval zebrafish leads naturally to future directions that use this behavior as a model for investigating how homeostatic and allostatic control systems are implemented by the brain. This endeavor would leverage the unique advantages - genetic and optical accessibility [38,39] - of the larval zebrafish for uncovering the neural bases of behavior. Such studies would also dovetail nicely with previous work on the neural circuitry underlying phototaxis [10,13,15]. It would also be particularly interesting to identify where in the brain the luminance set point is computed and stored. The results of our study suggest that the set point computation requires luminance information from both eyes, so the computation likely occurs at a site in the brain that receives visual input from both eyes. One candidate brain region is the torus longitudinalis, which receives visual information from both eyes [40,41], responds to changes in luminance with sustained firing [42], and plays a role in orienting behaviors towards light [43]. A description of the set point’s neural implementation will also allow targeted studies on how its value is updated when environmental luminance changes. Because homeostatic and allostatic control systems play vital roles in many aspects of animal physiology and behavior, understanding the implementation and evolution of the set point for luminance-based navigation could also yield insights into other physiological control systems.

## Acknowledgements

We thank the members of the Engert lab and Ahrens lab for general feedback, as well as the following individuals for detailed feedback on and editing of the manuscript: Kristian J. Herrera, Yasuko Isoe, Robert E. Johnson, Lucy Lai, Stephen C. Thornquist, Hanna Zwaka.

A.B. was supported by the Human Frontier Science Program Long-Term Fellowship LT000626/2016. F.E. received funding from the National Institutes of Health (U19NS104653, R43OD024879, 2R44OD024879), the National Science Foundation (IIS-1912293) and the Simons Foundation (SCGB 542973).

This material is based upon work supported by the National Science Foundation Graduate Research Fellowship Program awarded to A.B.C. under Grant No. DGE1745303. Any opinions, findings, and conclusions or recommendations expressed in this material are those of the authors and do not necessarily reflect the views of the National Science Foundation.

## Author Contributions

Conceptualization: A.B.C. and F.E.; Methodology: A.B.C.; Software: A.B.C., D.D., and A.B.; Formal Analysis: A.B.C.; Investigation: A.B.C. and D.D.; Data Curation: A.B.C. and D.D.; Writing - Original Draft: A.B.C., D.D., and F.E.; Writing - Review and Editing: A.B.C., D.D., and F.E.; Supervision: A.B.C. and F.E.; Funding Acquisition: F.E.

## Declaration of Interests

The authors declare no competing interests.

## METHODS

### LEAD CONTACT AND MATERIALS AVAILABILITY

#### Data and Code Availability

The swim data and analysis code used to generate figures 1 - 3 are available on GitHub (https://github.com/diptodip/brightfish/experiments). Raw frame data is available upon request. The code used in the modeling experiments for Figure 4 is also available on GitHub (https://github.com/diptodip/brightfish).

#### Materials Availability

This study did not generate new unique reagents.

### EXPERIMENTAL MODEL AND SUBJECT DETAILS

#### Zebrafish

For all experiments, we used wild-type larval zebrafish (strains AB and WIK), aged 5 to 8 days post-fertilization (dpf). We did not determine the sexes of fish we used. Fish were raised in shallow Petri dishes and fed *ad libitum* with paramecia after 4 dpf. Fish were raised on a 14/10 light/dark cycle at around 27C. All experiments were done during daylight hours (4 - 14 hours after lights on). All protocols and procedures were approved by the Harvard University/Faculty of Arts and Sciences Standing Committee on the Use of Animals in Research and Teaching (Institutional Animal Care and Use Committee).

### METHOD DETAILS

#### Design of system for tracking and closed-loop video projection

For behavioral experiments related to Figure 1 - 3, we used the same behavioral system for tracking freely swimming larval zebrafish as in [18]. Larval zebrafish swam freely in custom-made, circular, acrylic dishes with black walls and filled with filtered system water. Dish diameter was 12 cm; wall height was 5 mm. Fish were bottom-lit using light-emitting diode (LED) arrays (940 nm, Cop Security) so that the shadow they cast could be used to determine their position and orientation. We tracked the fish using a Grasshopper3-NIR camera (FLIR Systems) equipped with a zoom lens (Zoom 7000, 18-108mm, Navitar) and a long-pass filter (R72, Hoya). Frame data were stably acquired at around 90 frames per second and analyzed in real time to extract fish position and orientation. For each experiment, we used 2 groups of 4 cameras, with each group of 4 connected to a different computer; thus, each computer could track 4 separate fish simultaneously. Fish were tracked using code written in C++, Python 3.7, and OpenCV 4.1 (See Quantification and Statistical Analyses). We delivered visual stimuli locked to the position and orientation of the fish.

#### Visual Stimuli

We delivered different luminance values to the fish by commanding a video projector (60 Hz, AAXA P300 Pico Projector) to project different grayscale pixel values, ranging from 0 (black) to 1 (white). We used an iPhone 11 Pro and the Lux Light Meter Pro app at a distance of about 5 cm above the dish to measure brightness levels: 0 < 1 Lux; 0.25 = 240 Lux; 0.5 = 345 Lux; 0.75 = 425 Lux; 1 = 560 Lux (Supplemental Figure 1A). Visual stimuli were projected from below onto white paper to disperse the light for visibility. Visual stimuli were projected in the reference frame of the fish. For split-luminance experiments, all pixels with negative x values in this coordinate system were defined to be left of the fish, and all pixels with positive x values were defined to be right of the fish.

#### Set point seeking experiments

Related to Figure 1. We held the fish in either dim (L = 0.25) or bright (L = 0.75) luminance for 10 seconds, and then subjected a fish to a split-luminance test period in which the luminance of the left and right sides of the fish were selected randomly from five possible pixel values (0, 0.25, 0.5, 0.75, 1). For the bright preadaptation experiments, the test period lasted for 10 seconds; however, we noticed a decrease in turning bias after the first few seconds due to adaptation (Supplemental Figure 1C-D). As a result, we limited the test period to 3 seconds for the dim preadaptation experiments and only analyzed the first 3 seconds of the test period in both conditions.

#### Split-luminance preadaptation experiments

Related to Figure 2. We held the fish in a split luminance environment (luminance on dim side: L = 0; luminance on bright side: L = 1) for 16 seconds. After this preadaptation period, we changed the luminance of either the bright side (for Figure 2) or the dim side (for Supplemental Figure 2E-G) for 10 seconds.

#### Two-preadaptation experiments

Related to Figure 3. We held the fish in an initial preadaptation period (L = 0.25 or L = 0.75) for 16 seconds. We then held the fish in a second preadaptation period; if the luminance during the initial preadaptation was L = 0.25, the luminance during the second preadaptation was L = 0.75. On the other hand, if the luminance during the initial preadaptation was L = 0.75, the luminance during the second preadaptation was L = 0.25. Across different trials, the length of the second preadaptation period was chosen pseudorandomly from these possible values: 0 s (i.e. no second preadaptation), 3 s, 6 s, 9 s, and 12 s.

#### Modeling

Complete details on our *in silico* experiment settings and initializations of fish parameters are available on GitHub at https://github.com/diptodip/brightfish.

We simulate two computational models of zebrafish phototaxis -- one using separate monocular information and one integrating binocular information. All simulations occur within a 2D grid of dimensions (H, W) (rows, columns). In our simulations, both H, W = 101 for the spotlight experiment (Figure 4G) and H, W = 51 for the partitioned halves experiments (Figure 2, Figure 4D). We simulate fish as points without volume within this grid. For both models, the fish calculates brightness in each eye as the mean value of all grid tiles falling within two coterminal rays originating from the fish position with an angle of 0.8π between them.

On each time step, the simulated fish first updates its set point(s). The monocular fish has two set points, one for the left eye (**SP**_**L**_) and one for the right eye (**SP**_**R**_). It will calculate the differences

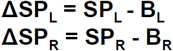

And then use a learning rate **r** to update its set points:

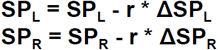

The binocular fish has one set point. It will calculate the difference

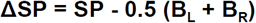

And similarly use a learning rate to update the set point:

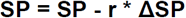

After updating its set point(s), the fish turns in the direction of the eye with a smaller difference from the set point as described in Results; if **E**_**L**_ **- E**_**R**_ **> 0**, the fish turns right and otherwise the fish turns left. The fish samples two turn angles from two Normal distributions — one from a no turn distribution **Δ θ**_**n**_ **∼ N(0**.**01, 0**.**50)** and the other from the turn distribution **Δ θ**_**t**_ **∼ N(0**.**52, 0**.**59)**. The sign of the turn direction is flipped for right turns versus left turns and we describe angles in radians. These distributions were generated by fitting Gaussian curves to turn angle distributions of the preadaptation period (no turn distribution) or test period (turn distribution) of the condition in Figure 1 where preadaptation luminance was 0.75 and the test period luminances were (0.75,0) (data not shown but are available on GitHub). The fish calculates a linear combination **Δ θ = (1 - λ)(Δ θ**_**n**_**) + (λ Δ θ**_**t**_), where **λ = f(E**_**L**_ **- E**_**R**_) and **f** is a nonlinear function **f(x) =** |**x**|^**(1/3)**^, **x** ∈ **[-1, 1]**. The choice of exponent does not greatly affect fish behavior (data not shown). Then, the fish updates its heading as θ = (θ + Δ θ) mod 2π. Finally, the fish determines whether a swim occurs on this time step by sampling from a Bernoulli distribution with probability **p**_**move**_. If the fish swims on this time step, it samples a move distance **d ∼ N(μ**_**d**_, **σ**_**d**_) and moves **d** units in its heading direction **θ**.

### QUANTIFICATION AND STATISTICAL ANALYSIS

#### Closed-loop tracking and swim-bout detection for freely swimming zebrafish

Software used for tracking freely swimming larval zebrafish and detecting swim bouts in real time was the same as in [18]. First, the image background was calculated as the mode image over approximately 60 seconds. For each acquired image frame, the background was subtracted. In the mode-subtracted image, the center of mass was defined to be the position of the larval zebrafish, and we used second-order image moments to determine its orientation. To detect swim bouts, we calculated a variance over a rolling, 50 ms time window. Variance spikes, subject to interbout interval constraints, were used to determine swim bout times.

#### Statistical Tests

Details of statistical tests used in this study can be found in the figure legends. Unless otherwise specified in the figure legends, error bars signify +/- standard deviation around the mean. Trials were selected pseudorandomly using the output of a random number generator and without human input. Sample sizes were not predetermined. We excluded fish that appeared for long periods in the image background, as this suggested that they were dead or otherwise immobile. All exclusion was done prior to data analysis.

## SUPPLEMENTAL INFORMATION

**Supplemental Figure 1.**
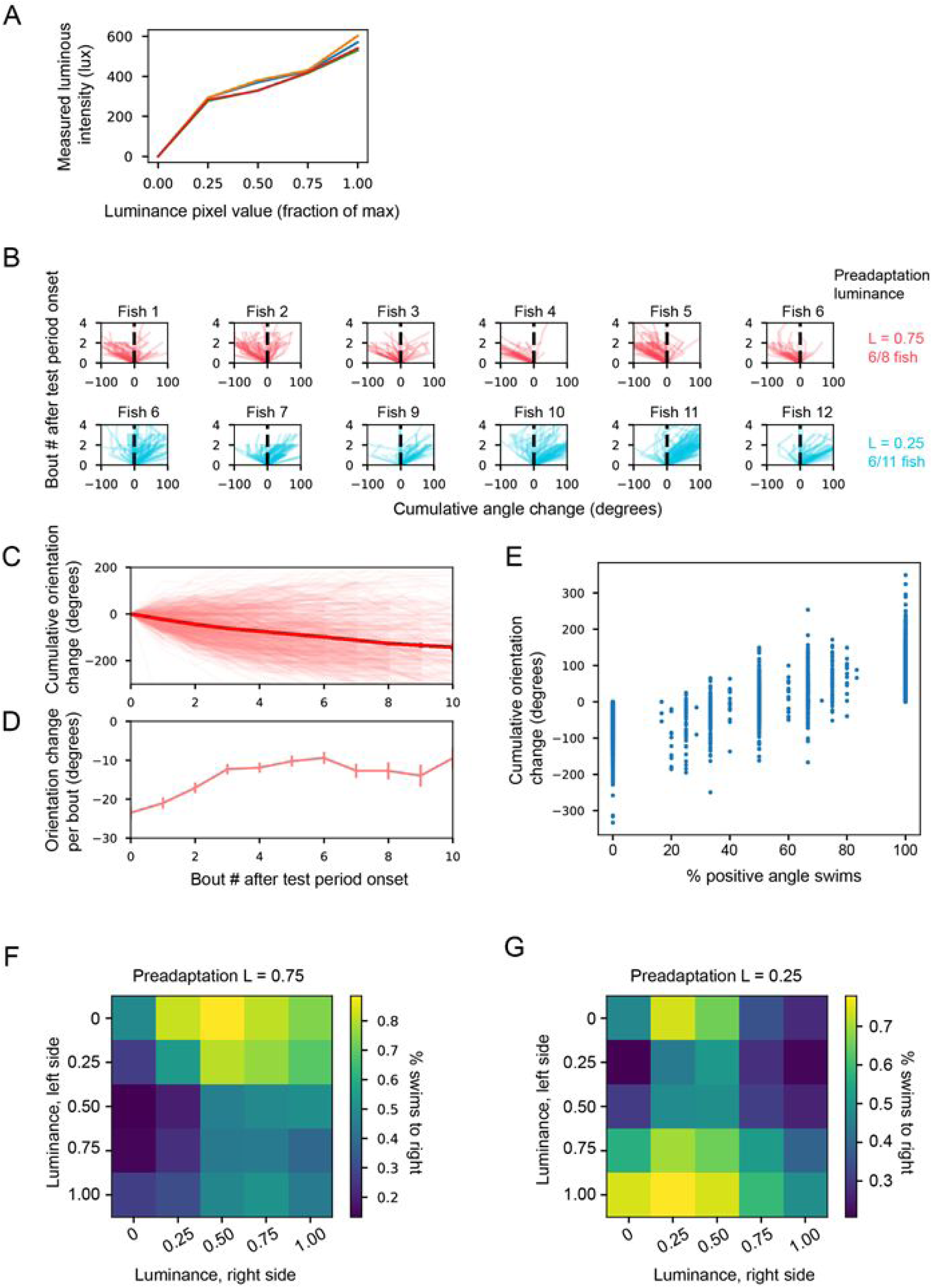
related to Figure 1. **A**. Calibration curves of 4 projectors used for visual projection during behavioral experiments. Measured illumination intensity (in lux) is plotted against pixel value (as a fraction of maximum). **B**. Single fish cumulative turning angles in the first 3 seconds of the split luminance period. Each graph corresponds to an individual trace. Each trace corresponds to cumulative angle change in a trial. Top row, red traces: preadaptation luminance is 0.75. Bottom row, blue traces: preadaptation luminance is 0.25. Data from the first 6 of 8 (preadaptation 0.75) and first 6 of 11 (preadaptation 0.25) fish are presented here. **C**. Cumulative angle changes across all trials and fish over the entire 10 s test period (preadaptation luminance: 0.75, test period left luminance left: 1, test period luminance right: 0). This corresponds to a longer test period time window for the red traces shown in Figure 1E. **D**. Mean and standard deviation of per-bout angle change for data shown in C. The magnitude of angle change for the first 3 bouts is significantly greater than that for later bouts (p < 0.05, t-test). **E**. Relationship between % of swims to the right and cumulative angle turn, demonstrating good correspondence between the cumulative angle metrics used in Figure 1C-F and the % swims metric used in Figure 1G-I (linear relationship, slope: 116.75 degrees, intercept: -56.92 degrees, r: 0.74, t-test on slope p < 10^−6^. **F-G**. Mean % turns to the left (defined to be negative angle turns) for different test period combinations of left and right side luminances. Panel E corresponds to a preadaptation luminance of 0.75, while panel F corresponds to a preadapation luminance of 0.25. Note that these are the same data plotted against E_R_ - E_L_ in Figure 1I.

**Supplemental Figure 2.**
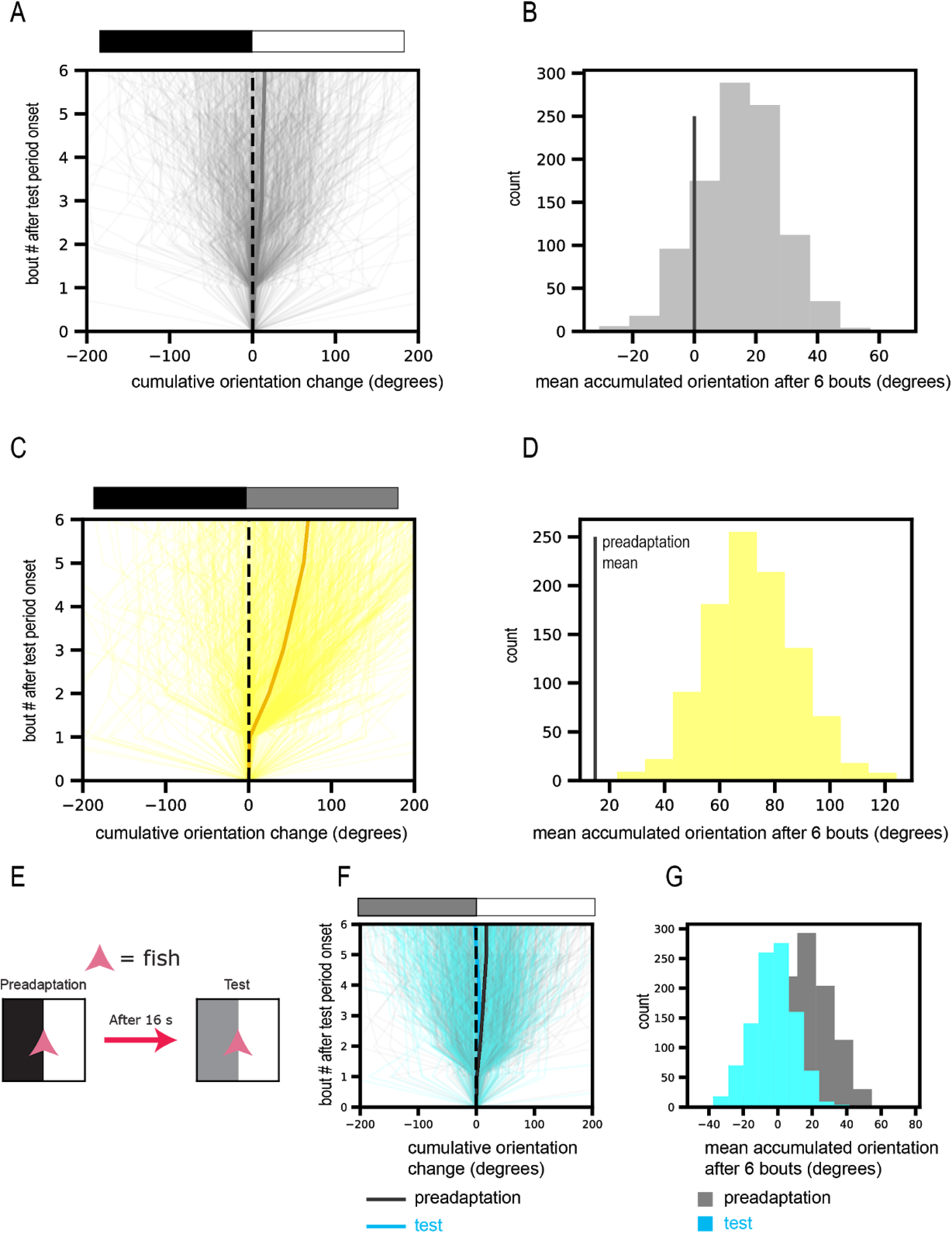
related to Figure 2. **A**. Cumulative turning angle during the last 5 seconds of the preadaptation period before onset of the test period. Each thin trace depicts data from one trial. Bold gray line shows the mean cumulative angle. **B**. Bootstrapped distribution of the mean cumulative angle after 6 swim bouts for the data presented in A. Vertical line denotes 0 degrees. The mean of the cumulative angle is not significantly different from zero (75% CI [-0.70 degrees, 29.47 degrees]). Mean of 50 sampled trials, nBoot = 1000. **C**. Same as A, except for the first five seconds of the test period. Note that these data are presented in Figure 2I but are reproduced here for comparison. Thick yellow line shows the mean cumulative angle. **D**. Bootstrapped distribution of the mean cumulative angle after 6 swim bouts for the data presented in C. Vertical line denotes 0 degrees. The mean of the cumulative angle is significantly larger than 0 degrees (99% CI [30.40 degrees, 117.77 degrees]. Mean of 50 sampled trials, nBoot = 1000. **F**. Cumulative turning angles for experiments shown in G. Thin gray traces - last 5 seconds of preadaptation period. Thin blue traces - first 5 seconds of test period. Thick blue and gray traces show the mean cumulative angles for test and preadaptation periods, respectively. **G**. Bootstrapped mean cumulative turning angles for the test (blue) and preadaptation (gray) period data shown in F. The distributions are significantly different (2 sample t test, p < 10^−10^). Mean of 50 sampled trials, nBoot = 1000

## Notes

https://github.com/diptodip/brightfish

### Competing Interest Statement

The authors have declared no competing interest.

